# Subjective optimality in finite sequential decision-making

**DOI:** 10.1101/2020.07.15.204321

**Authors:** Yeonju Shin, HeeYoung Seon, Yun Kyoung Shin, Oh-Sang Kwon, Dongil Chung

**Author notes:** These authors contributed equally to this work (Yeonju Shin, HeeYoung Seon). Correspondence should be made to O.-S.K. or D.C.

## Abstract

Many decisions in life are sequential and constrained by a time window. Although mathematically derived optimal solutions exist, it has been reported that humans often deviate from making optimal choices. Here, we used a secretary problem, a classic example of finite sequential decision-making, and investigated the mechanisms underlying individuals’ suboptimal choices. Across three independent experiments, we found that a dynamic programming model comprising subjective value function explains individuals’ deviations from optimality and predicts the choice behaviors under fewer opportunities. We further identified that pupil dilation reflected the levels of decision difficulty and subsequent choices to accept or reject the stimulus at each opportunity. The value sensitivity, a model-based estimate that characterizes each individual’s subjective valuation, correlated with the extent to which individuals’ physiological responses tracked stimuli information. Our results provide model-based and physiological evidence for subjective valuation in finite sequential decision-making, rediscovering human suboptimality in subjectively optimal decision-making processes.

## Introduction

Hiring a new employee is one of the toughest decisions to make as a team leader. Most of the time, there are only a limited number of job openings available and a limited time period in which to complete the hiring process. This process is even more difficult when applicants are accepted on a rolling basis, because one has to make a choice whether to accept the current applicant without knowing whether other future potential applicants would have been a better fit for the job. Likewise, there are many decision problems in life that are sequential and constrained by a certain time window. The ‘secretary problem’ is a classic example of this finite sequential decision problem and has been widely used to understand the optimal policy in making choices (e.g., to hire or not) under a limited number of opportunities (Ferguson, 1989; Freeman, 1983). Provided with the full information (i.e., the distribution of candidates), the optimal solution for the problem is to choose the first number that is above a mathematically calculated decision threshold (Hill & Krengel, 1991). However, it is not clear whether and how humans deviate from optimal choices. Here, we used one variant of the secretary problem, in which the distribution of candidates is given and the reward is the value of the chosen candidate, to investigate (i) whether individuals make the optimal decision in a finite sequential decision problem, and (ii) if not, how do they make their decisions. Our results provide behavioral and physiological evidence supporting that individuals make threshold-based choices in a finite sequential decision problem and that seemingly suboptimal decision patterns (deviation from the optimal) originate from the process of optimally calculating thresholds using individuals’ subjective value function.

To examine how individuals make choices in a finite sequential decision problem, we recorded behavioral choices, response time (RT), and pupil dilation of 91 participants (male/female = 45/46, age = 22.88 ± 1.93 years) as they made a series of choices to accept or reject a random number presented on the screen (**Fig. 1A**). During each round, they had a fixed amount of opportunities (chances) to evaluate a new random number by rejecting previously presented numbers. When they accepted, the presented number was added to their final payoff, and then they moved on to the next round (up to 200 rounds) that consisted of a new set of chances. Overall, we implemented three separate experiments. In Experiment 1, participants had up to five opportunities (K = 5), and they were not explicitly informed of the maximum number that would be presented. Experiment 2 had up to two or five opportunities (K = 2 or 5), and the participants were informed of the full distribution information (including the maximum). In Experiment 3, to temporally dissociate actions (choosing to accept or reject) from physiological responses to stimuli, participants were not allowed to make a choice until an audio cue was played. All the other settings were equal to Experiment 2 where participants had up to five chances (K = 5). Non-overlapping samples were obtained from each experiment (see **Materials and Methods** for detailed experimental procedures).

**Figure 1.**
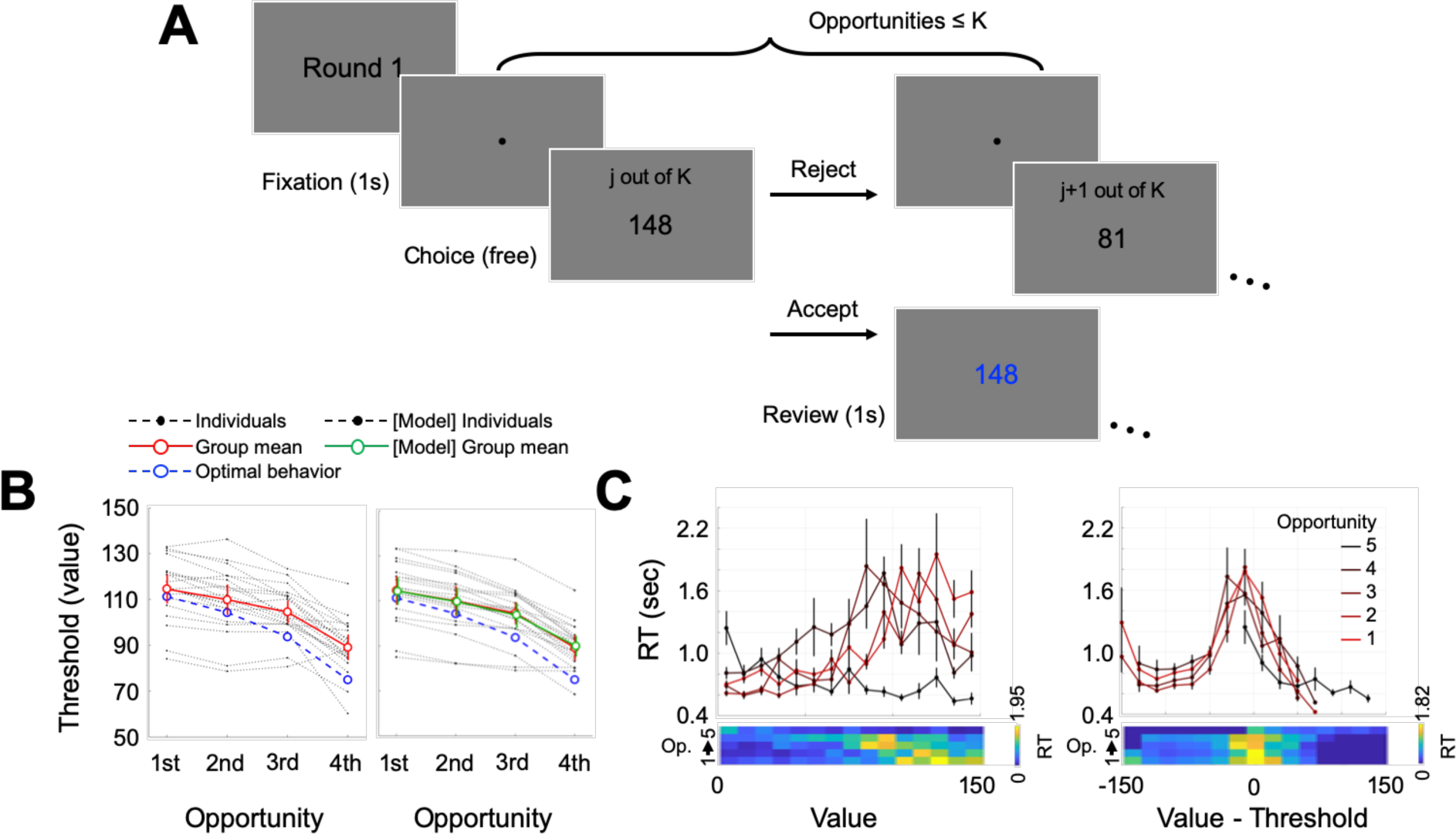
Experimental procedures and behavioral results of Experiment 1. **(A)** Participants made a series of choices between accepting and rejecting a presented number. At each round, they had up to K opportunities (K = 5 in Experiment 1, K = 2 or 5 in Experiment 2) to reject the number and get a new random number; the round ended when participants accepted a presented number. At the last opportunity, participants were given no choice but to accept the presented number. A new set of stimuli (numbers) was used in the next round. **(B)** The optimal decision threshold per opportunity (blue), calculated under the assumption of the full information, was compared with a corresponding empirical decision threshold (red). **(C)** Response times (RTs) for each opportunity were computed against the presented stimuli values. Regardless of the opportunity, RTs showed negative association with the absolute distance between the presented stimuli and the corresponding decision threshold. That is, participants showed the shortest RTs for the numbers that are farthest from decision thresholds, and vice versa. Error bars represent s.e.m.

## Results

### Experiment 1

#### Individuals show higher decision thresholds than the optimal decision model

Each presented number, sampled from a uniform distribution ranging from 0 to 150, could be considered as an option whose value matches its face value (the number). Because individuals can only accept a single number within each round, they should accept a number only when it is large enough. Specifically, an optimal decision-maker should not accept a presented number unless it is larger than the expected value of successive opportunities. For example, individuals should accept any numbers at the last opportunity (i.e., the fifth opportunity in Experiment 1) and thus the expected value of the last opportunity is 75. Based on this information, at the opportunity one before the last (the fourth in Experiment 1), a value-maximizing individual should accept any numbers higher than 75 but reject other numbers. Following the dynamic programming approach,(Bellman, 1966) we computed an optimal threshold for each opportunity (**Fig. 1B**, blue).

To examine whether individuals follow such decision processes, we calculated empirical thresholds—the value where individuals were equally likely to accept or reject—from 20 participants’ behavioral choices (male/female = 10/10, age = 22.85 ± 1.31 years) (**Fig. 1B**). Consistent with the optimal thresholds (blue), empirical thresholds (red) at the later opportunities were lower than those at the earlier opportunities (mean threshold differences between the first and the second = 4.70, t(19) = 5.24, Cohen’s d = 1.17, *p* = 4.66e-5; the second and the third = 5.39, t(19) = 3.99, Cohen’s d = 0.89, *p* = 7.84e-4; and the third and the fourth = 15.38, t(19) = 7.10, Cohen’s d = 1.59, *p* = 9.41e-7). However, participants showed empirical thresholds significantly higher than the optimal thresholds, indicating that people have higher expectations about later opportunities (the difference between empirical and optimal thresholds = 8.49, t(19) = 3.28, Cohen’s d = 0.73, *p* = 0.004).

Compared with optimal thresholds, it is not difficult to notice that the average empirical thresholds have a shallower slope as evidenced by the increasing difference between the empirical and optimal thresholds across opportunities (mean slope of [empirical - optimal]: 3.75, t(19) = 4.40, Cohen’s d = 0.98, *p* = 3.08e-4). Although the empirical decision thresholds suggest otherwise, one may still suspect that an alternative heuristic individuals might have used was to apply a constant threshold regardless of the number of remaining opportunities (i.e., applying a constant threshold across all opportunities). It is well known that easier choices—here, deciding whether to accept or not the presented value that is far smaller or larger than the threshold—require shorter response times (RT) (Ratcliff, 1978). If individuals applied the same threshold across all opportunities, mean RTs should be symmetric around a certain value (i.e., threshold). To examine this possibility, we calculated mean RT within each opportunity. The symmetric pattern was observed only when mean RTs were calculated as a function of presented values adjusting for the estimated empirical threshold within each corresponding opportunity (**Fig. 1C**). This result suggests that individuals did apply differential thresholds for each opportunity during decision-making.

#### Subjective optimality explains individual choice patterns

Prospect theory has suggested that outcomes are perceived as gains and losses relative to a certain reference point, and that gains and losses are valued following concave and convex subjective value functions, respectively (Tversky & Kahneman, 1979). We drew on this framework to evaluate potential decision processes accounting for individuals’ sub-optimal decision thresholds. In accordance with Prospect theory (Tversky & Kahneman, 1979), we hypothesized that individuals’ subjective valuation (U) for a given value (*v*) is dependent on their individual reference point (r) and nonlinear value sensitivity (ρ), as follows:

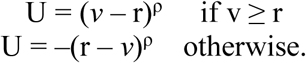

Note, we focused on valuation *per se*, and thus, the time it took for individuals to establish (learn) their reference points (their own perspective of the environment) was assumed negligible (see **Discussion** for further consideration of learning effects). Importantly, two additional components were introduced. First, individuals may perceive the waiting time till acceptance costly and take it into account in valuation. Second, we hypothesized that this subjective value-based computation occurs not only during active decision-making, but also at mental simulation such that individuals use their subjective valuation in constructing expectations of each opportunity (i.e., computing decision thresholds; **Fig. 2A**).

**Figure 2.**
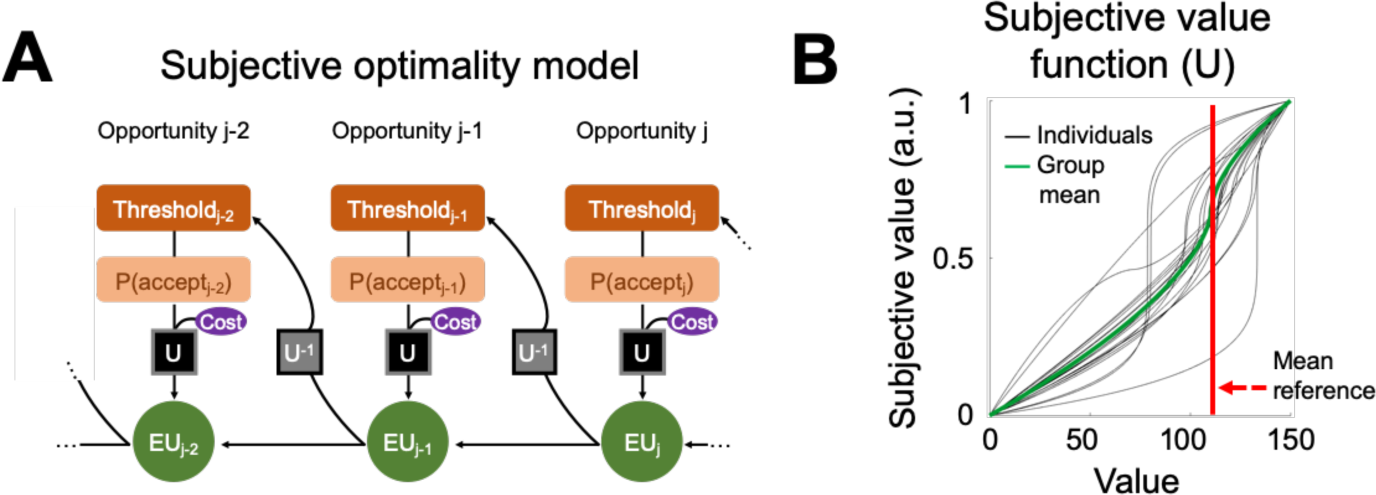
Subjective optimality model. **(A)** The optimal decision model assumes that individuals compute the decision threshold of a certain opportunity based on the expected value of successive opportunities. In the ‘Subjective optimality model’, expected values of the successive opportunities are replaced by expected utilities (EU) calculated based on the subjective value function as per Prospective theory. **(B)** Two free parameters, reference point, and nonlinear value sensitivity define subjective valuation of the presented stimuli values. Group average subjective value function (green) is depicted using the group mean of individual estimates: reference point = 114.77; value sensitivity = 0.47.

This ‘Subjective optimality model’ with a waiting cost converges to three nested models in special cases: the Subjective optimality model without a waiting cost (Cost = 0), the Optimal decision model (ρ = 1), and the Constant threshold model (ρ = 0) (see **Materials and Methods** for model details). A formal model comparison using a likelihood ratio test revealed that the Subjective optimality models with and without a potential waiting cost explained individuals’ choice behaviors comparably well (*χ*^2^(20) = 19, *p* = 0.52). Moreover, these models showed superior explanatory power compared to the two other nested decision models (**Table S1**). These results suggest that the waiting cost was negligible in Experiment 1, values larger than the reference point (114.77; **Fig. 2B**) were perceived as gains, and any value stimuli smaller than the reference point were perceived as potential losses. Moreover, this result indicates that individuals use marginally diminishing (concave) and increasing (convex) subjective value function for gains and losses, respectively, in finite sequential decision-making.

### Experiment 2

#### The Subjective optimality model predicts behavioral alterations in the context of scarce opportunity

In our suggested model, change of reference point reframes one’s subjective valuation and, in turn, alters decision thresholds. Given this causal relationship, we can predict that one would lower their decision threshold in the context where one expects less overall outcome and consequently sets a lower reference point. To examine whether empirical data matches the prediction from the model, we conducted a second experiment where some participants had five (K = 5) and other participants had two opportunities (K = 2) in each round (**Fig. 1A**). That is, in contrast to Experiment 1, individuals who had two opportunities always had to accept the second value if they rejected the first presented stimulus. If individuals followed the Optimal decision model, the decision threshold at the first opportunity among K = 2 should be equal to the decision threshold at the fourth opportunity among K = 5. As predicted from the Subjective optimality model, the decision thresholds estimated from participants (K = 2: N = 23, male/female = 11/12, age = 23.09 ± 2.09 years; K = 5: N = 21, male/female = 11/10, age = 23.19 ± 1.86 years; non-overlapping from Experiment 1) were significantly different depending on the number of opportunities one had per round (threshold_K=5, 4th_ = 90.42 ± 9.72, threshold_K=2, 1st_ = 79.28 ± 15.45; t(42) = 2.83, Cohen’s d = 0.85, *p* = 0.007; **Fig. 3A**). Of note, different from Experiment 1, the Subjective optimality model better explained participants’ empirical choices for K = 5 when a waiting cost was included as a free parameter (**Table S1**). However, the group mean of the estimated waiting cost was not different from zero (t(20) = 0.37, Cohen’s d = 0.08, *p* = 0.71), suggesting that the additional parameter was needed to explain individual differences in their subjective waiting costs.

**Figure 3.**
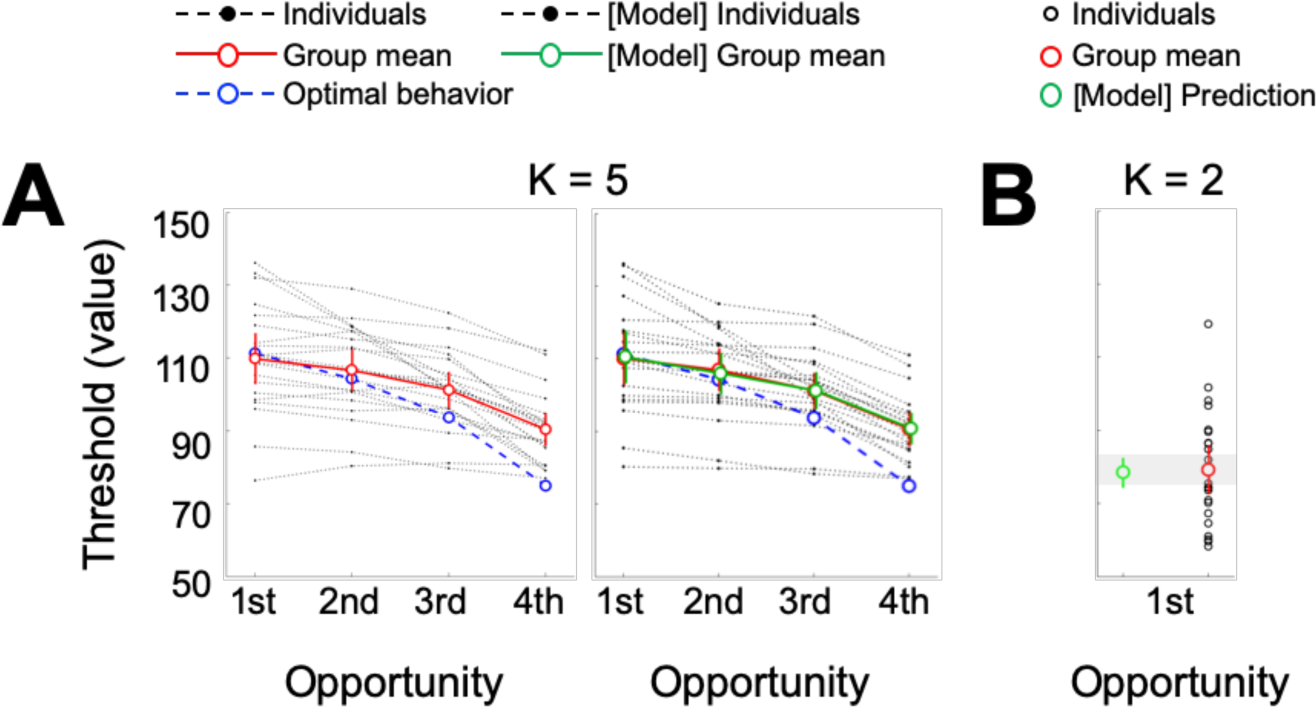
Behavioral results of Experiment 2. **(A)** In individuals who had five opportunities (K = 5), empirical decision thresholds (red) along the opportunities were comparable with that of Experiment 1. To examine whether or not our Subjective optimality model can be generalized to other contexts, a model prediction of the decision threshold was made for K = 2 (green); value sensitivity was assumed to be the same even in the different context, but the reference point was set to a lower level adjusted proportionately to the reduction of the expected payoff. **(B)** Empirical (observed) decision threshold in individuals who had two opportunities (K = 2) was consistent with the prediction. Error bars represent s.e.m.

Next, we examined whether our model quantitatively captures behavioral alterations dependent on the scarcity of opportunities. By lowering the reference point parameter proportionately to the extent of expected payoff reduction and keeping all the other parameters the same, the model-based threshold prediction for K = 2 (78.50 ± 4.09; **Fig. 3B**, green) was consistent with the observed behavioral threshold (see **Materials and Methods** for model prediction details), which supports the critical role of the reference point in subjective valuation. One may suggest that the task with K = 2 is simple enough for participants and that they would have followed the optimal strategy (i.e., using 75 as a decision threshold at the first opportunity). However, this is unlikely given that only 7 out of 23 participants’ credible intervals of the empirical decision threshold, defined by the 95% highest density interval, included 75 (see **Materials and Methods**). Furthermore, the large across-individual variability in behavioral decision thresholds (SD = 15.45; **Fig. 3B**) showcased that the Optimal decision model cannot explain individuals’ decision strategies. These results again support the Subjective optimality model suggesting that individuals make threshold-based choices in a finite sequential decision problem, and that seemingly suboptimal decision patterns (e.g., waiting for future chances) may have originated from the process of calculating thresholds using individuals’ subjective value function.

### Experiment 3

To further investigate physiological instantiation of the decision processes implemented in our model, we examined changes of pupil diameter acquired while participants made a series of choices. A rich set of evidence suggests that pupil dilation (or contraction) reflects not only individuals’ arousal level (Nassar et al., 2012; Urai, Braun, & Donner, 2017), but also cognitively complex information, such as value (Van Slooten, Jahfari, Knapen, & Theeuwes, 2018), uncertainty (Urai et al., 2017), cognitive conflict (Cavanagh, Wiecki, Kochar, & Frank, 2014), and choice (de Gee, Knapen, & Donner, 2014). Drawing upon these findings, we hypothesized that, should subjective valuation occur as proposed, changes of pupil diameter may capture value of stimuli, decision difficulties, and final choices that participants would make. To test this hypothesis, we operated a slightly modified task; (i) participants had to view the presented stimuli for a period of time (1.5-2.5 seconds) before being allowed to accept or reject the stimuli, and (ii) an audio cue was used to announce to participants that they could make a choice (**Fig. 4A**). This modification temporally dissociated choice from other cognitive processes (e.g., valuation) and prevented the introduction of any visual confounds in analyzing physiological signals at the time of decision-making. Participants had up to five opportunities per each round, and all other experimental settings were equal to Experiment 2 (see **Materials and Methods** for details).

**Figure 4.**
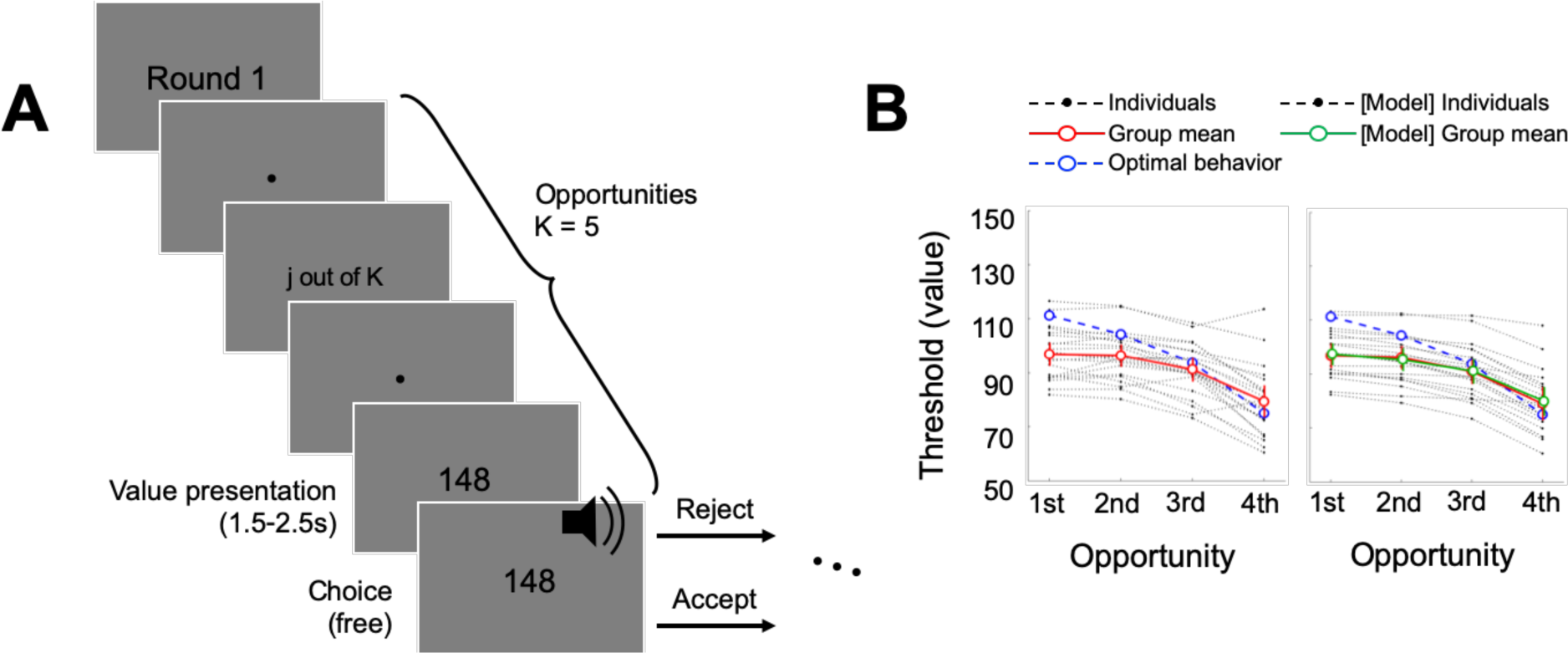
Experimental procedures and behavioral results of Experiment 3. **(A)** To temporally dissociate valuation from action selection, we implemented a modified task design where individuals had to wait for an audio cue to make choices. **(B)** Empirical decision thresholds (red) were compared with the optimal decision thresholds (blue). Compared with Experiments 1 and 2, in Experiment 3, individuals showed lower decision thresholds at the early opportunities. Error bars represent s.e.m.

### Waiting is costly

Twenty-two new participants were recruited for Experiment 3 (10 females, age = 22.59 ± 2.32 years; non-overlapping from Experiments 1 or 2). With the addition of forced waiting time, which accumulated over opportunities, we expected that participants would perceive choices to accept after a longer wait less valuable (Kable & Glimcher, 2007; Loewenstein & Prelec, 1992) and thus, they would accept earlier. Consistent with our expectation, a stark difference in the behavioral pattern was observed in Experiment 3 compared to Experiments 1 and 2. Specifically, decision thresholds from empirical data in Experiment 3 (red solid line) were below the optimal decision thresholds (blue dotted line), indicating that participants were more likely to accept small numbers that they would have rejected in the other two experimental settings (**Fig. 4B**).

This result was corroborated by the model-based results. First, the Subjective optimality model with a waiting cost showed superior explanatory power for Experiment 3 compared with alternative models (**Table S1**), emphasizing again that the waiting cost plays an important role in finite sequential decision-making. Second, the average of the estimated waiting cost parameter was significantly larger than zero only in Experiment 3 (t(20) = 63.51, Cohen’s d = 13.86, *p* = 1.51e-24), and it was larger than the cost parameters in the other two experiments (Experiment 3 > 1: t(40) = 15.67, Cohen’s d = 4.84, *p* < 1.00e-15; Experiment 3 > 2: t(41) = 5.64, Cohen’s d = 1.72, *p* =1.41e-6; **Fig. 5**). Third, as it was intended from the task modification, individuals’ behavioral change was sourced specifically back to the waiting cost parameter, such that other parameters (nonlinear value sensitivity and reference point) were not affected (**Fig. 5**). These results together support our interpretation suggesting that the perceived cost of waiting underlies the behavioral alteration in the new task environment.

**Figure 5.**
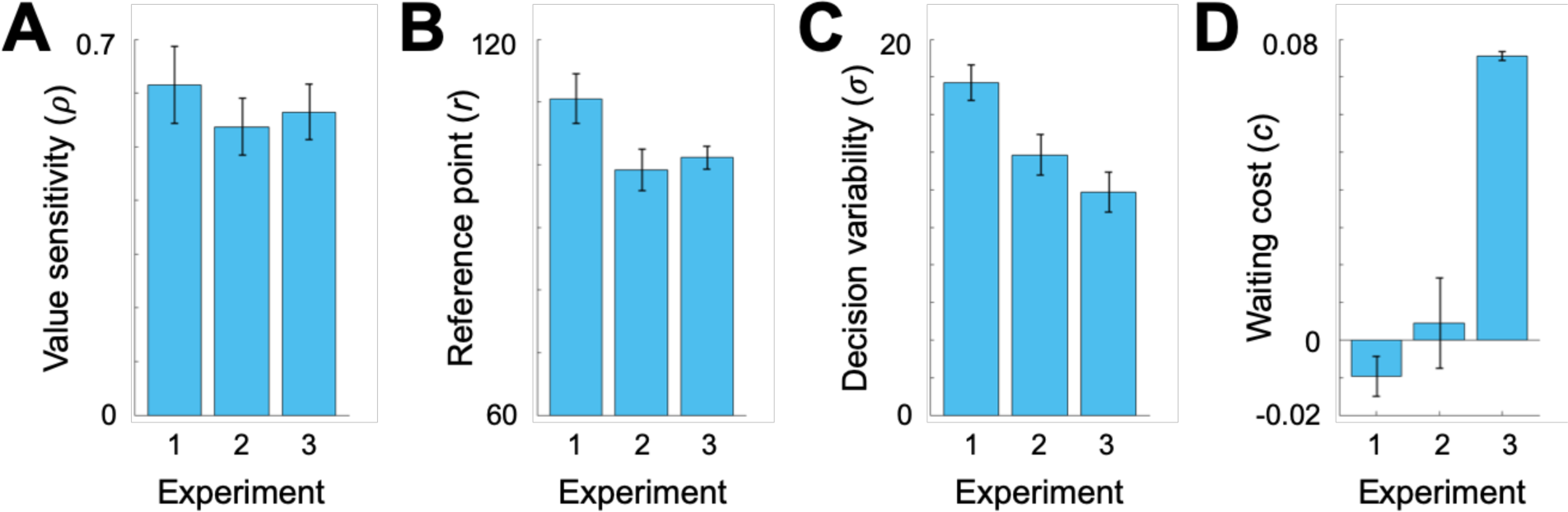
Best fitting parameters. The Subjective optimality model was used to estimate the four parameters that explain individuals’ behavioral choices. **(A)** The estimated nonlinear value sensitivity (ρ) was comparable among all three separate experiments (Experiments 1, 2 (K = 5), and 3: F(2, 59) = 0.45, *p* = 0.64). **(B)** There was a significant difference in reference points between experiments (F(2, 59) = 3.67, *p* = 0.032). Post-hoc tests revealed that the difference originates from the higher reference point in Experiment 1 where participants were not informed of the maximum stimuli value (Tukey test: Experiment 1 vs. 2: *p* = 0.036; Experiment 1 vs. 3: *p* = 0.099; Experiment 2 vs. 3: *p* = 0.893). **(C)** There was a significant difference in decision variability between experiments (F(2, 59) = 8.00, *p* = 8.40e-4). Post-hoc tests revealed that the difference originates from the higher decision variability in Experiment 1 (Tukey test: Experiment 1 vs. 2: *p* = 0.031; Experiment 1 vs. 3: *p* = 0.001; Experiment 2 vs. 3: *p* = 0.372). **(D)** Waiting costs were larger than zero only in Experiment 3 (Experiment 1: t(19) = -1.84, Cohen’s d = -0.41, *p* = 0.082; Experiment 2: t(20) = 0.37, Cohen’s d = 0.08, *p* = 0.71; Experiment 3: t(20) = 63.51, Cohen’s d = 13.86, *p* = 1.51e-24). Moreover, the estimated waiting cost in Experiment 3 was significantly larger than those in the other two Experiments (Experiment 3 > 1: t(40) = 15.67, Cohen’s d = 4.84, *p* < 1.00e-15; Experiment 3 > 2: t(41) = 5.64, Cohen’s d = 1.72, *p* = 1.41e-6). Error bars indicate s.e.m.

### Pupil dilation reflects choice and decision difficulty

As described above, we then examined whether physiological responses reflect cognitive decision processes. First, we compared pupil diameter changes between accepted and rejected opportunities. Consistent with previous reports, pupil size was significantly different depending on the subsequent choices (de Gee et al., 2014) (**Fig. 6A**). Particularly, pupil dilations within 558-726 msec and 1182-1500 msec were associated with subsequent acceptance of the presented values (t(17) > 2.11, all *p*s < 0.05). Only the latter cluster remained significant after controlling for multiple comparisons using a cluster-based permutation method (Maris & Oostenveld, 2007) (numerical *p* = 3.50e-4). Still, given the fact that the time of the earlier cluster (558-726 msec) overlaps with the range of RTs in Experiments 1 and 2 (**Fig. 1C, S1**), this result suggests that participants may have covertly made choices as early as 550 msec and the cognitive process was reflected in the physiological responses (de Gee et al., 2014) (see **Fig. S2** for a pupil size result reflecting individuals’ arousal level).

**Figure 6.**
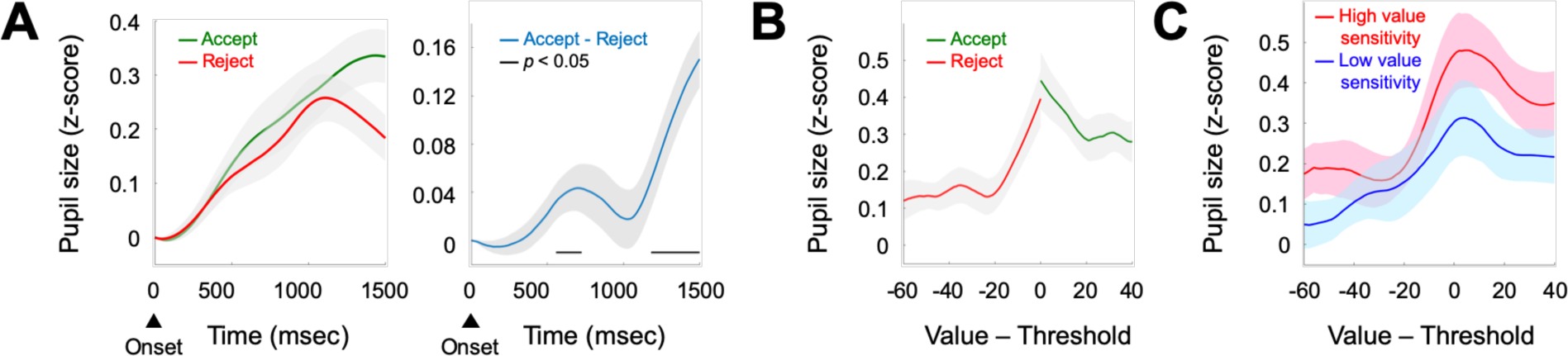
Pupillometry responses reflect subsequent choices and decision values. **(A)** Pupil size change from the stimuli onset was measured, separately for the accepted (green) and rejected (red) opportunities. Paired comparison between the cases revealed significant pupil dilation for the accepted stimuli at the early stage after the onset, and again at the later time. **(B)** To examine whether or not pupil size reflected stimuli value, pupil size 1500 msec after the stimuli onset was depicted as a function of the signed distance between stimuli value and the corresponding decision threshold. **(C)** Individuals who had higher value sensitivity in their estimated parameter (median split; red) showed more pronounced pupillometric responses reflecting the value information. Shades represent s.e.m.

Next, we calculated mean pupil diameters as a function of subjective values. This was done for accepted and rejected stimuli separately, so that the relationship between pupil sizes and values is independent of subsequent choices. Regardless of the choice, as we observed from RT patterns (**Fig. 1C, S1**), pupil size was negatively correlated with ‘decision difficulty’. That is, in both rejected and accepted trials, pupil size decreased as a function of the absolute distance between the decision threshold and value of the presented stimuli (Rejected trials: slope = -0.0043, t(17) = - 2.48, Cohen’s d = -0.58, *p* = 0.024; Accepted trials: slope = -0.0039, t(17) = -2.49, Cohen’s d = - 0.59, *p* = 0.024; **Fig. 6B**). Steepness of the slopes was comparable between accepted and rejected opportunities (t(17) = 0.19, Cohen’s d = 0.05, *p* = 0.85). However, the intercept, i.e., pupil dilation at the corresponding threshold, was higher for accepted than rejected trials (t(17) = 2.13, Cohen’s d = 0.50, *p* = 0.046). Furthermore, pupil sizes between accepted and rejected trials were significantly different even after controlling for the distance between stimuli and the threshold (t(17) = 3.62, Cohen’s d = 0.85, *p* = 0.002), which indicates that pupil sizes reflect additional information other than decision difficulty. Together, these results suggest that pupil dilation reflects both decision difficulty and subsequent choices (Cavanagh et al., 2014; de Gee et al., 2014), the two crucial components comprising subjective valuation (Kolling et al., 2016; Rangel, Camerer, & Montague, 2008).

### Physiological sensitivity matches behavioral value sensitivity

As evidenced by the model parameter estimates, there are individual differences in the extent to which one responds to a unit increase of presented stimulus value (i.e., value sensitivity). We tested whether or not this modeling construct of individual characteristics matches with individuals’ physiological responses. To provide an illustrative description, we divided participants into two subgroups based on their parameter estimation (median split) where one group had lower value sensitivity and the other group had higher value sensitivity. For each group, we calculated average pupil dilation as a function of signed decision difficulty (the difference between stimulus value and the decision threshold of the corresponding opportunity) (**Fig. 6C)**. Individuals who had high value sensitivity (red) showed relatively high pupil dilation compared to individuals who had low value sensitivity (blue). This positive correlation between value sensitivity and pupil dilation was statistically significant at the threshold where decision difficulty is the highest (Pearson’s correlation r = 0.52, *p* = 0.027). The result indicates that individuals who have high behavioral value sensitivity indeed have higher physiological sensitivity to stimuli value. Moreover, the consistent patterns across physiological and behavioral data reflecting individuals’ characteristics serve as additional evidence suggesting the use of subjective valuation in finite sequential decision-making.

## Discussion

Our results provide a model-based explanation for suboptimality in finite sequential decision-making. Specifically, we present evidence that subjective valuation reflecting individuals’ belief about the environment underlies the mechanism of how the brain computes decision thresholds in the problem.

As a classic example of a finite sequential decision problem, various versions of the secretary problem were investigated (Ferguson, 1989; Freeman, 1983). The standard secretary problem simulates the cases where only the relative ranks matter, such that individuals have to find the best option (e.g., a candidate in a hiring scenario) among the sequentially presented options (Chow, Moriguti, Robbins, & Samuels, 1964; Guan & Lee, 2018). In this setting, inferior choices (choosing options that are not the best) lead to no reward, but we have to note that this is hardly the case in real-life. First, any choices we make should have some value even in the case where they were not the best option. For example, an employee who ends up not meeting the employer’s original expectation still can make some contribution (except for an unfortunate case in which the employee turns out to be a con artist and shuts down the business). Second, in reality, it is impossible for the decision maker to learn the true relative rank of the chosen option, because the decision maker will have no knowledge about the subsequent options that were to follow. In other words, there is no one who can examine the success of the choice and deliver a reward if, and only if, the choice were correct. The current study addressed this discrepancy by implementing a task where each option had a monetary reward that matched its face value. Although there was no explicit instruction saying that individuals should find the best option within the finite number of opportunities, participants were informed that the final payoff would be determined by the accumulated reward amount across the entire task and thus, the task preserved the goal of reward maximization. We believe that the current variation of the secretary problem provides a more naturalistic setting to investigate individuals’ sequential decision-making.

A typical behavior pattern observed across various versions of the secretary problem is that individuals show suboptimal choices, such that they wait less than the optimal stopping point (Bearden, Rapoport, & Murphy, 2006; Seale & Rapoport, 1997). This suboptimal choice tendency is accounted for by lower decision thresholds than the optimal decision threshold, indicating that they are more likely to accept the option that has low value. Our results across the three experiments may seem inconsistent from this perspective. Particularly, individuals showed higher thresholds for both Experiments 1 and 2, but lower thresholds for Experiment 3. The main change in Experiment 3 was the additional forced wait introduced before the cue when participants were allowed to submit their choice. Our model-based analysis results suggest that this subtle change in task design may have triggered participants to think more about the tradeoff between payoffs and time they spent per round. Such an impact of additional ‘cost of waiting (extra time)’ is consistent with previous reports showing that non-zero interview cost was associated with lowering decision thresholds (Costa & Averbeck, 2015; Seale & Rapoport, 1997; Yeo, 1998). Our model parameter estimates supported this interpretation, such that only in Experiment 3, the estimated cost was significantly larger than zero. These results highlight that the context of decision-making (e.g., task schedule) as well as the extent to which individuals find the task costly (e.g., cognitively demanding or mentally boring) are crucial in decision-making processes (Kool, McGuire, Rosen, & Botvinick, 2010).

Our Subjective optimality model included two free parameters essential in capturing individuals’ choice patterns. First, the reference point reflects each individual’s belief about the environment (Tversky & Kahneman, 1979). It is known that beliefs can alter how individuals respond to given information, which not only affects their behavioral choices, but also neural responses (Gu et al., 2015). In line with this, we showed that discouraged expectation (scarce opportunities in Experiment 2) causes individuals to be more pessimistic about future chances and wait less in deciding (lowering thresholds). In addition, interestingly, individuals’ expectations (reference point) were significantly higher when they did not have full information about stimuli distribution (Experiment 1). This result suggests that humans, in general, have optimistic bias (Sharot, Korn, & Dolan, 2011), which may diminish or even become inverted in other contexts (e.g., scarce opportunities, mental costs).

Second, the nonlinear value sensitivity indicates the extent to which individuals’ subjective valuation increases for an additional unit of reward. In the current study, the sensitivity represented as an exponent term in the utility function was smaller than one, which captures marginally diminishing returns for gains and marginally increasing returns for losses (Tversky & Kahneman, 1979). In our suggested model, a range of value sensitivity characterizes a spectrum of decision characteristics in individuals. Value sensitivity close to zero represents a rather categorical valuation (gain or loss relative to the reference point) and choices that are accounted for by a constant threshold being insensitive to the context (i.e., remaining opportunities). On the other hand, value sensitivity close to one represents objective valuation and choices that follow the Optimal decision model. In concert with the reference point, individuals’ value sensitivity shapes the extent to which they take into account uncertainty of future opportunities in decision-making. This wide range of individual differences may explain why some individuals are more stubborn with their opinions (e.g., stereotype), while others easily adapt to contextual information (Taylor, 1981).

In the current study, the pupil responses encode both decision difficulty and the subsequent choice of whether individuals will accept or reject the presented stimulus. Both types of information temporally preceded actual choice, so these pupil dilations are the physiological representations of the processed information regarding decision-making, rather than a simple reflection of the presented visual information. As suggested from previous studies, pupil dilation may reflect the downstream processing of the anterior cingulate cortex (Cavanagh et al., 2014; Critchley, Tang, Glaser, Butterworth, & Dolan, 2005), the brain region that is involved in encoding decision difficulty (Shenhav, Straccia, Cohen, & Botvinick, 2014), and, more broadly, a wealth of value-related information—including difficulty signals—during decision-making processes (Kolling et al., 2016). Differential pupil sizes depending on the subsequent choices suggest that there is more to neurophysiological representation than simple decision difficulties. Individuals may pay more attention to the stimuli that they plan to accept for accumulating more evidence (Krajbich, Armel, & Rangel, 2010). Of course, such a process may have the opposite causality, in that the rich amount of accumulated evidence of a particular stimulus may induce even higher attention levels (e.g., saliency driven bottom-up attention (Koch & Ullman, 1987)). In the current study, the latter is unlikely, given that all low-level visual information (e.g., contrast) of the displayed stimuli were matched or controlled for. The current results show that the two pieces of information essential in subjective valuation are linked together at the physiological level.

The future direction of the current study includes expanding our model to further explain the mechanisms of how individuals learn the stimulus distribution (e.g., reinforcement learning). In the current study, we assumed that the learning process is rapid and negligible in relevance to other decision processes. Previous studies reported no evidence of learning in various versions of the secretary problem (Campbell & Lee, 2006; Seale & Rapoport, 1997). Moreover, we showed that decision processes under imperfect information (no knowledge of the maximum stimuli value) were comparable with the processes under the full information. This result suggests that, even without explicit information about the stimuli distribution, people, in general, have a rough idea about the range of values of an uncertain option. Alternatively, people were able to learn early enough (Goldstein, McAfee, Suri, & Wright, 2020) that the behavioral strategy for the rest of the task was not different from the case where individuals knew about the distribution from the beginning. Still, inclusion of learning mechanisms in the model would be essential to examine whether or not the decision model generalizes to broader contexts (e.g., a volatile environment). Examples of finite sequential decision problems span a wide range of life choices, including finding the right life partner and choosing a career, the aims of which are to maximize reward under a limited amount of resources and opportunities. Such value-based decision processes with reference to costs are not unique to humans but extend from fish choosing a mate, who become less selective under costly environments (Milinski & Bakker, 1992), to primates making foraging decisions (Hayden, Pearson, & Platt, 2011). The Subjective optimality model provides a way in which individual subjective valuation generates systematic biases in sequential decision-making and opens a window to decompose physiological responses into decision difficulty and signatures of subsequent choice, of which levels differ in the extent of individual value sensitivity. In sum, our data support a mechanistic account of suboptimal choices varying from overly impulsive choices in individuals with substance-use problems (Ekhtiari, Victor, & Paulus, 2017) to delayed choices in individuals who suffer from indecisiveness (Rassin & Muris, 2005).

## Materials and Methods

### Participants

Ninety-one healthy young adults (male/female = 45/46, age = 22.88 ± 1.93 years) participated in the current study. All participants provided written informed consent and were paid for their participation. The study was approved by the Institutional Review Board of Ulsan National Institution of Science and Technology (UNISTIRB-18-39-C, UNISTIRB-18-14-A). None of the participants reported a history of neurological or psychiatric illness. Three separate experiments were conducted and there were no overlapping participants across experiments. Twenty students participated in Experiment 1 (male/female = 10/10, age = 22.85 ± 1.31 years), and 47 students were recruited for Experiment 2 where they had five or two opportunities per round (male/female = 23/24, age = 23.00 ± 1.98 years). Among the participants in Experiment 2, three participants were excluded from the analyses due to their reported suspicion about the payment structure of the experiment. Among the included participants, 21 students (male/female = 11/10, age = 23.19 ± 1.86 years) were randomly assigned to the condition where they were given five opportunities per round, and 23 participants (male/female = 11/12, age = 23.09 ± 2.09 years) were assigned to the condition where they were given two opportunities per round. Twenty-four students participated in Experiment 3 (male/female = 12/12, age = 22.67 ± 2.28 years). Two participants were excluded due to their reported suspicion about the payment structure of the experiment, and one participant was excluded due to data loss from a computer error. Three participants were excluded from the pupil diameter analyses due to poor calibration. After exclusion, data from 21 participants (male/female = 11/10, age = 22.62 ± 2.38 years) were used for behavioral analyses, and a subsample of the data (N = 18; male/female = 8/10, age = 22.33 ± 2.30 years) was used for further pupil diameter analyses. All participants reported normal or corrected-to-normal vision under soft contact lenses (no glasses were allowed due to potential reflections during eye-tracking).

### Stimuli and apparatus

All stimuli were generated using Psychophysics Toolbox Version 3 (www.psychtoolbox.org) and MATLAB R2017a (MathWorks), and presented on a DLP projector (PROPixx VPX-PRO-5050B; screen size of 163 × 92 cm^2^; resolution of 1920 × 1080 pixels; refresh rate of 120 Hz; linear gamma). The distance between the participants’ eyes and screen was fixed at 153 cm. The ambient and background luminance were set at 1.1 and 69.2 cd/m^2^, respectively. The main stimuli were three-digit integer numbers, randomly selected between zero and 150. To minimize luminance effects on pupil size, one-or two-digit numbers were displayed as three-digit numbers with extra zeros attached in front of the stimuli (e.g., 1 is displayed as ‘001’). During the task, fixation was enforced at the center of the screen with an infrared eye tracker (Eyelink 1000 Plus, SR Research, Canada), and a chin and forehead rest were used to minimize head movement.

### Experiments

At the beginning of the task, the eye-tracker was calibrated, referencing eye fixation data at the four corners of the screen. During the task, participants made a series of choices either to accept or to reject presented stimuli (**Fig. 1A**). As explained above, the stimuli were randomly selected integers between zero and 150 where each number had equal probability of being selected (uniform distribution). Each presented number could be considered as an option whose value matches its face value because participants were instructed that all accepted numbers would be added to their final payoff at the end of the task. Given this knowledge, participants had a fixed number of opportunities (chances) to evaluate and reject a new randomly selected number. The present ‘round’ ended when participants accepted a presented number within this limited number of opportunities, or when they ran out of the opportunities where they had no other choice but to accept the presented number at the last chance. At the beginning of each opportunity, participants were shown which opportunity they were currently at, so that they would not lose track of the number of remaining opportunities. A new round followed, at which the number of available opportunities was reset to the original maximum quantity. Participants were paid at the end of the study (after completing 200 rounds), based on the sum of the numbers they chose during the task. All instructions were provided through illustrated slides.

Overall, we implemented three separate experiments, each of which had slightly different settings. In Experiment 1, participants had up to five opportunities (K = 5), and they were not explicitly informed of the maximum number (150) that would be presented. Use of the context with incomplete information was to incorporate a more naturalistic setting as real-life problems where, as in most of the cases, individuals do not have knowledge about the best potential option (e.g., even if the current candidate for a job has a good enough fit for the position, one cannot assure that a potential future candidate will not have a superior fit). Participants were instructed that the presented stimuli would be sampled from a uniform distribution, and thus, we expected that participants would quickly deduce the maximum range through iterative experiences. At the beginning of the new round, the accumulated payoff amount up until the last round was presented at the bottom of the screen. In Experiment 2, participants were randomly assigned to one of two conditions where one condition had five (K = 5) and one condition had two (K = 2) opportunities. Here, participants were also informed of the maximum number (i.e., 150). In addition, participants were given a practice session that comprised two rounds where all the stimuli were ‘000’, which allowed them to be familiarized with associated buttons and the task screen settings. All the rest of the task settings were identical to Experiment 1.

Experiment 3 was designed to temporally dissociate actions (i.e., accept or reject) from the stimulus onset, so that physiological responses to stimuli independent from potential motor preparatory signals could be measured. Particularly in Experiment 3, participants were not allowed to make choices until an audio cue was played (**Fig. 4**). The audio cue was played between 1.5 and 2.5 seconds after stimulus onset (uniform distribution), which allowed us to tease out potential confounding factors related to action from the pupil diameter measures at 0-1.5 seconds after stimulus onset. In addition, to prevent participants from making unnecessary eye movements, all the information including number stimuli were presented at the center of the screen. As implemented in Experiment 2 where K = 5, participants were informed that the maximum number was 150 and that they have up to five chances to evaluate the stimuli per each round.

### Behavioral analysis

For all three tasks, behavioral choices (accept or reject) and response time (RT) were measured. Individuals’ decision threshold for each opportunity was estimated from their choices. To estimate empirical decision threshold for each opportunity, a cumulative distribution function of Gaussian distribution was fitted to individuals’ choice data that corresponded to the same opportunity across all 200 rounds. The mean and variance parameters of the Gaussian distribution represent the decision threshold and decision variability, respectively. A set of best-fitting parameters that maximize the likelihood of the data was estimated per individual using the Nelder-Mead simplex algorithm provided by MATLAB R2017b.

### Computational modeling and model comparison

For a formal model comparison at the group level, choices from all 200 rounds per participant were used for parameter estimation. We used likelihood-ratio tests to compare goodness-of-fit of the models for explaining participants’ decisions.

### Optimal decision model

An optimal decision maker is expected to maximize their payoff by estimating the expected value of each opportunity. This computation can be conducted from the final opportunity to the first, given the full information about the stimuli distribution (U[0, 150]). For example, in a condition where K = 5, the expected value of the last opportunity is 75, and therefore a payoff-maximizing optimal decision maker should set 75 as the decision threshold of the fourth opportunity (i.e., accept numbers larger than 75 and reject those that are lower). Then, this decision strategy should again determine the expected value of the fourth opportunity. Generalizing this dynamic programming approach, the decision threshold of the i^th^ opportunity (*ϑ*[i]) can be written as follows:

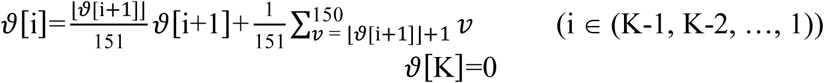

where ⌊ *x*⌋ indicates the greatest integer less than or equal to *x*.

### Subjective optimality model

Our hypothesis was that individuals use subjective valuation in reference to their own expectations about the environment during finite sequential decision-making. To test the hypothesis, we constructed a computational model drawn upon Prospect theory (Kahneman & Tversky, 1979). Particularly, individuals’ subjective valuation (U) of an objective value (*v*) was defined as below:

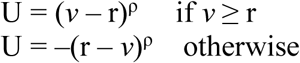

where ρ and r indicate individuals’ nonlinear value sensitivity and reference point, respectively. Subjective valuation is also used in computing decision thresholds:

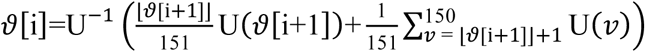

where U^-1^(.) indicates an inverse function of the aforementioned subjective value function.

### Subjective optimality model with a waiting cost

In our secretary problem task, choosing to reject the current stimulus means that participants have to go through further steps (opportunities) to receive rewards (or at least to find out how much reward they will receive) until they choose to accept at a later opportunity. Such an additional wait may introduce a disutility (i.e., negative value) against the choice to reject. To test this possibility and quantitatively estimate this ‘mental waiting cost’, we modified our suggested Subjective optimality model to a more general format as follows:

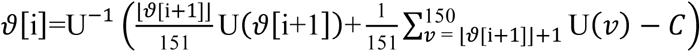

where *C* indicates a waiting cost per opportunity. Note that the waiting cost lowers the expected utility of the following opportunity (i+1^th^), and thus has an effect of lowering the decision threshold of the current opportunity (i^th^).

### Constant threshold model

There is a simple alternative decision strategy for the secretary problem: to use a constant decision threshold throughout all opportunities. To examine this possibility, we estimated one decision threshold per individual. This constant threshold model provides a quantitative baseline for a formal model comparison.

### Predicting change of decision threshold based on the altered decision context

To examine whether or not our suggested model can be generalized under different contexts with scarce opportunities, we took a prediction approach using model-based information from the context with abundant opportunities. Specifically, the reference point and nonlinear value sensitivity parameters estimated from behavioral choices of individuals (N=21) who participated in Experiment 2, K = 5 were used to predict the decision threshold in the two opportunities condition (K = 2). Particularly for the nonlinear value sensitivity, the parameter distribution in the K = 2 condition was assumed to be the same as that in the K = 5 condition. On the other hand, the parameter distribution of the reference point was assumed to be shifted down by the difference of expected earnings between the two conditions, reflecting participants’ acknowledgement of the scarce number of opportunities. To be agnostic about differential subjective valuation under different contexts, the change of participants’ expectation about mean earning was calculated comparing expected values between conditions. To predict the mean threshold in the K = 2 condition, 23 pairs of parameters (matching the number of participants in K = 2) were randomly sampled with replacement from the aforementioned parameter distribution, and the thresholds corresponding to each parameter pair were computed by applying our model. The procedure was repeated 5,000 times to estimate the distribution of the mean of 23 thresholds. The 95% confidence interval was computed from the 5,000 means.

### Parameter estimation procedure

We used Bayesian hierarchical analysis to estimate the best-fitting parameters for participants’ choice data (Daw, 2011). The parameters characterizing individual participants were drawn from the population distributions, each of which follows a Gaussian distribution. The priors on the means of the population distributions (µ) were set to broad uniform distributions, and the priors on the SDs (σ) were set to an inverse-Gamma distribution in each of which, the shape parameter alpha is one and the scale parameter beta is manually selected. To improve sampling efficiency, we sampled the parameters from a transformed space, and the hierarchical structure was assumed in the transformed space. Specifically, the reference point and value sensitivity parameters were sampled without domain restrictions and transformed by a scaled logistic function *g*(*x*) = A/(1+exp(-*x*)) before applying to the model. In the function *g*(*x*), A was set to 150 for the reference point parameter r, and set to 2 for the values sensitivity parameter ρ. The decision variability parameters and the group-level hyper-parameters for parameters’ standard deviation were transformed by exp(.) after sampling. We did not apply a transformation to the waiting cost parameter. A Markov chain Monte Carlo (MCMC) method (Metropolis-Hastings algorithm) was used to sample from the posterior density of the parameters conditioned on all of the participants’ choices. We estimated the most likely set of parameters for each participant from the resulting chain of samples using a multivariate Gaussian kernel function provided by MATLAB R2017b.

### Pupillometry: Preprocessing

Pupil diameter was sampled at 500 Hz from both eyes using an infrared eye-tracker (Eyelink 1000 Plus; SR Research, Kanata, Canada) and recorded continuously for the entire session. Blinks and saccades in each eye were identified using the standard criteria provided by Eyelink, and the identified intervals were linearly interpolated. Particularly for the blink events, the interpolation was applied to the intervals between 150 ms before and after each identified blink. Three participants whose pupil data included a large proportion of interpolated intervals (> 50 %) were excluded from further analyses. The means of the interpolated data from both eyes were band-pass filtered between 0.02-4 Hz using third-order Butterworth filters. The long-lasting effects (∼ 5 sec) of blinks on pupil diameter were identified by applying least-squares deconvolution to individual data, and then removed from the data (Knapen et al., 2016). Then, the resulting data were z-scored for each session (i.e., each participant). Pupil diameter changes in response to the value stimulus were computed for each opportunity. Each epoch was defined for pupil responses between -200 and 1,500 msec around the stimulus onset, and corrected for its baseline by subtracting the mean pupil size around (± 20 msec) the onset. The choice trials that required a large proportion (> 50%) of interpolation were excluded from the analysis, which comprised 28% of the entire choice trials.

### Pupillometry: Statistical tests

To examine whether physiological responses reflect cognitive decision processes, we tested pupil dilations and contractions in response to (i) subsequent choices to accept or reject, and (ii) decision difficulty. First, pupil diameter changes between 0-1,500 msec after the stimulus onset were compared between accepted and rejected opportunities. We used t-tests to compare mean differences at each time step and defined statistically significant temporal clusters (alpha level set to 0.05). To control for the false alarm rate, we used the cluster-based permutation method (Maris & Oostenveld, 2007) and examined the statistical significance of each cluster. Particularly in the permutation procedure, the sign of the difference value for each participant was randomized and the sum of t-values in each cluster was used as its statistic. Second, the pupil dilation at 1,500 msec after the stimulus onset was used to examine the effect of decision difficulty—the absolute distance between the corresponding decision threshold and the presented value—on the pupil dilation. Linear regression was used for the rejected trials (choice = reject, - 40 < value - threshold < 5) and accepted trials (choice = accept, -5 < value - threshold < 40) separately for each participant. The same set of data points was used to test the effect of choice on pupil dilation after controlling for the decision difficulty. We further investigated individual differences in the extent to which one responds to stimulus value at the physiological level (i.e., pupil dilation). Pupil dilation at 1500 msec after the stimulus onset was used. We smoothed each individual’s pupil dilation data along the threshold centered values from -90 to 60 by applying local regression using a 2D polynomial model provided by MATLAB R2017b. The estimated pupil dilation at threshold was used to calculate the Pearson correlation between individuals’ estimated value sensitivity and their pupil responses.

## Author contributions

Y.K.S., D.C., and O.-S.K. designed the experiments. Y.S., H.S., D.C., and O.-S.K. analyzed the data. H.S., D.C. and O.-S.K. drafted the manuscript. All of the authors discussed the results, revised, and approved the final manuscript.

## Data and materials availability

The datasets generated during and/or analyzed during the current study are available from the corresponding authors on reasonable request.

## References

Bearden, J. N., Rapoport, A., & Murphy, R. O. (2006). Experimental studies of sequential selection and assignment with relative ranks. Journal of Behavioral Decision Making, 19(3), 229–250.

Bellman, R. (1966). Dynamic programming. Science, 153(3731), 34–37.

Campbell, J., & Lee, M. D. (2006). The Effect of Feedback and Financial Reward on Human Performance Solving’Secretary’Problems. Paper presented at the Proceedings of the Annual Meeting of the Cognitive Science Society.

Cavanagh, J. F., Wiecki, T. V., Kochar, A., & Frank, M. J. (2014). Eye tracking and pupillometry are indicators of dissociable latent decision processes. Journal of Experimental Psychology: General, 143(4), 1476.

Chow, Y., Moriguti, S., Robbins, H., & Samuels, S. (1964). Optimal selection based on relative rank (the “secretary problem”). Israel Journal of mathematics, 2(2), 81–90.

Costa, V. D., & Averbeck, B. B. (2015). Frontal–parietal and limbic-striatal activity underlies information sampling in the best choice problem. Cerebral cortex, 25(4), 972–982.

Critchley, H. D., Tang, J., Glaser, D., Butterworth, B., & Dolan, R. J. (2005). Anterior cingulate activity during error and autonomic response. Neuroimage, 27(4), 885–895.

Daw, N. D. (2011). Trial-by-trial data analysis using computational models. Decision making, affect, and learning: Attention and performance XXIII, 23(1).

de Gee, J. W., Knapen, T., & Donner, T. H. (2014). Decision-related pupil dilation reflects upcoming choice and individual bias. Proceedings of the National Academy of Sciences, 111(5), E618–E625.

Ekhtiari, H., Victor, T. A., & Paulus, M. P. (2017). Aberrant decision-making and drug addiction—how strong is the evidence? Current opinion in behavioral sciences, 13, 25–33.

Ferguson, T. S. (1989). Who solved the secretary problem? Statistical science, 4(3), 282–289.

Freeman, P. (1983). The secretary problem and its extensions: A review. International Statistical Review/Revue Internationale de Statistique, 189–206.

Goldstein, D. G., McAfee, R. P., Suri, S., & Wright, J. R. (2020). Learning when to stop searching. Management Science, 66(3), 1375–1394.

Gu, X., Lohrenz, T., Salas, R., Baldwin, P. R., Soltani, A., Kirk, U., … Montague, P. R. (2015). Belief about nicotine selectively modulates value and reward prediction error signals in smokers. Proceedings of the National Academy of Sciences, 112(8), 2539–2544.

Guan, M., & Lee, M. D. (2018). The effect of goals and environments on human performance in optimal stopping problems. Decision, 5(4), 339.

Hayden, B. Y., Pearson, J. M., & Platt, M. L. (2011). Neuronal basis of sequential foraging decisions in a patchy environment. Nature neuroscience, 14(7), 933.

Hill, T. P., & Krengel, U. (1991). Minimax-optimal stop rules and distributions in secretary problems. The Annals of Probability, 19(1), 342–353.

Kable, J. W., & Glimcher, P. W. (2007). The neural correlates of subjective value during intertemporal choice. Nature neuroscience, 10(12), 1625–1633.

Kahneman, D., & Tversky, A. (1979). Prospect theory: An analysis of decision under risk. Econometrica: Journal of the Econometric Society, 263–291.

Knapen, T., de Gee, J. W., Brascamp, J., Nuiten, S., Hoppenbrouwers, S., & Theeuwes, J. (2016). Cognitive and ocular factors jointly determine pupil responses under equiluminance. PloS one, 11(5).

Koch, C., & Ullman, S. (1987). Shifts in selective visual attention: towards the underlying neural circuitry. In Matters of intelligence (pp. 115–141): Springer.

Kolling, N., Wittmann, M. K., Behrens, T. E., Boorman, E. D., Mars, R. B., & Rushworth, M. F. (2016). Value, search, persistence and model updating in anterior cingulate cortex. Nature neuroscience, 19(10), 1280.

Kool, W., McGuire, J. T., Rosen, Z. B., & Botvinick, M. M. (2010). Decision making and the avoidance of cognitive demand. Journal of Experimental Psychology: General, 139(4), 665.

Krajbich, I., Armel, C., & Rangel, A. (2010). Visual fixations and the computation and comparison of value in simple choice. Nature neuroscience, 13(10), 1292.

Loewenstein, G., & Prelec, D. (1992). Anomalies in intertemporal choice: Evidence and an interpretation. The Quarterly Journal of Economics, 107(2), 573–597.

Maris, E., & Oostenveld, R. (2007). Nonparametric statistical testing of EEG-and MEG-data. Journal of neuroscience methods, 164(1), 177–190.

Milinski, M., & Bakker, T. C. (1992). Costs influences sequential mate choice in sticklebacks, gasterosteus aculeatus. Proceedings of the Royal Society of London. Series B: Biological Sciences, 250(1329), 229–233.

Nassar, M. R., Rumsey, K. M., Wilson, R. C., Parikh, K., Heasly, B., & Gold, J. I. (2012). Rational regulation of learning dynamics by pupil-linked arousal systems. Nature neuroscience, 15(7), 1040.

Rangel, A., Camerer, C., & Montague, P. R. (2008). A framework for studying the neurobiology of value-based decision making. Nature reviews neuroscience, 9(7), 545–556.

Rassin, E., & Muris, P. (2005). To be or not to be… indecisive: Gender differences, correlations with obsessive–compulsive complaints, and behavioural manifestation. Personality and Individual Differences, 38(5), 1175–1181.

Ratcliff, R. (1978). A theory of memory retrieval. Psychological review, 85(2), 59.

Seale, D. A., & Rapoport, A. (1997). Sequential decision making with relative ranks: An experimental investigation of the” secretary problem”. Organizational behavior and human decision processes, 69(3), 221–236.

Sharot, T., Korn, C. W., & Dolan, R. J. (2011). How unrealistic optimism is maintained in the face of reality. Nature neuroscience, 14(11), 1475.

Shenhav, A., Straccia, M. A., Cohen, J. D., & Botvinick, M. M. (2014). Anterior cingulate engagement in a foraging context reflects choice difficulty, not foraging value. Nature neuroscience, 17(9), 1249.

Taylor, S. E. (1981). A categorization approach to stereotyping. Cognitive processes in stereotyping and intergroup behavior, 832114.

Tversky, A., & Kahneman, D. (1979). Prospect theory: An analysis of decision under risk. Econometrica, 47(2), 263–291.

Urai, A. E., Braun, A., & Donner, T. H. (2017). Pupil-linked arousal is driven by decision uncertainty and alters serial choice bias. Nature communications, 8(1), 1–11.

Van Slooten, J. C., Jahfari, S., Knapen, T., & Theeuwes, J. (2018). How pupil responses track value-based decision-making during and after reinforcement learning. PLoS computational biology, 14(11), e1006632.

Yeo, G. F. (1998). Interview costs in the secretary problem. Australian & New Zealand Journal of Statistics, 40(2), 215–219.

