## Supplementary Information for "Subjective optimality in finite sequential decision-making"

**This PDF file includes:**

Figures S1 to S2
Tables S1

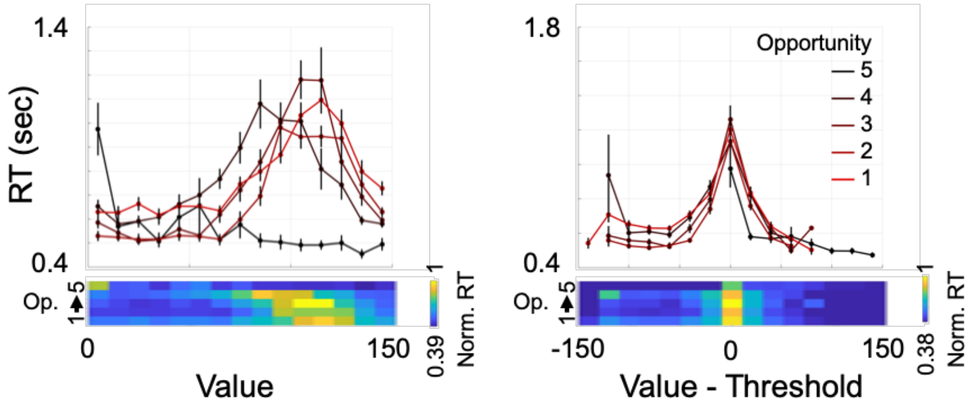


**Figure S1. Response times in Experiment 2**

Response times (RTs) for each opportunity in Experiment 2 were (Left) computed against the presented stimuli values. (Right) Regardless of the opportunity, RTs showed negative association with the absolute distance between the presented stimuli and the corresponding decision threshold. That is, participants showed the shortest RTs for the numbers that are farthest from decision thresholds, and vice versa. Error bars represent s.e.m.

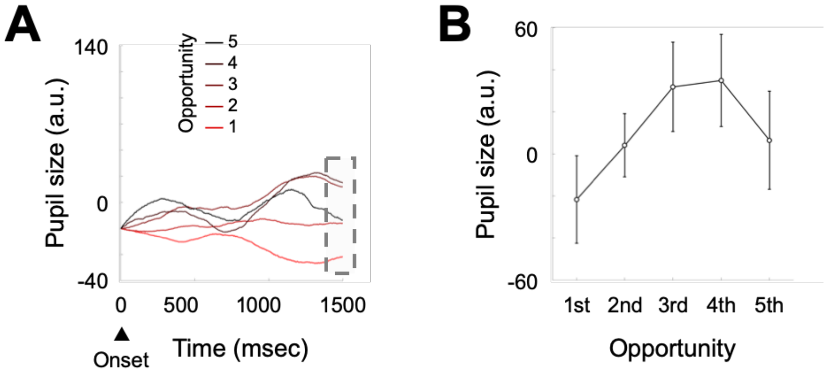


**Figure S2. Pupil responses reflect individuals' arousal level**

**(A)** Pupil size change from the stimuli onset was measured, separately for each opportunity ($1^{st}$, $2^{nd}$, …, and $5^{th}$). The pupil sizes at different opportunity were distinctively separable from around 1 sec after the stimuli onset. **(B)** Particularly, pupil sizes at 1500msec after the stimuli onset showed increasing pattern along the stages of opportunities, such that pupil size at the $1^{st}$ opportunity was significantly smaller compared with that at the $3^{rd}$ and $4^{th}$ opportunities (1st vs 3rd: $t(17) = 3.05$, Cohen's $d = 0.72$, $p = 0.007$; 1st vs 4th: $t(17) = 3.07$, Cohen's $d = 0.72$, $p = 0.007$). Such linearly increasing pattern in pupil responses along the repeated opportunities suggests that individuals entered higher arousal states as the number of remaining opportunities decreased. At the 5th opportunity, as the last opportunity in $K = 5$ condition, individuals had to accept any stimuli, which may explain why the pupil size at the 5th opportunity deviates from the linear pattern. Error bars represent s.e.m.

**Table S1.** Comparison of four models: Optimal decision model (Optimal), Constant threshold model (Constant), Subjective optimality model (SOptimal), and Subjective optimality model with waiting cost (SOptimal+C).

| | Constant | SOptimal | SOptimal+C |
|---|---|---|---|
| Experiment 1 | | | |
| Optimal | $\chi^2(20) = 846$<br>$p < 1.00e-15$ | $\chi^2(40) = 1408$<br>$p < 1.00e-15$ | $\chi^2(60) = 1427$<br>$p < 1.00e-15$ |
| Constant | | $\chi^2(20) = 561$<br>$p < 1.00e-15$ | $\chi^2(40) = 580$<br>$p < 1.00e-15$ |
| SOptimal | | | $\chi^2(20) = 19$<br>$p = 0.52$ |
| Experiment 2 | | | |
| Optimal | $\chi^2(21) = 1087$<br>$p < 1.00e-15$ | $\chi^2(42) = 1499$<br>$p < 1.00e-15$ | $\chi^2(63) = 1555$<br>$p < 1.00e-15$ |
| Constant | | $\chi^2(21) = 421$<br>$p < 1.00e-15$ | $\chi^2(42) = 468$<br>$p < 1.00e-15$ |
| SOptimal | | | $\chi^2(21) = 56$<br>$p = 5.58e-5$ |
| Experiment 3 | | | |
| Optimal | $\chi^2(21) = 1035$<br>$p < 1.00e-15$ | $\chi^2(42) = 1308$<br>$p < 1.00e-15$ | $\chi^2(63) = 1427$<br>$p < 1.00e-15$ |
| Constant | | $\chi^2(21) = 272$<br>$p < 1.00e-15$ | $\chi^2(42) = 391$<br>$p < 1.00e-15$ |
| SOptimal | | | $\chi^2(21) = 119$<br>$p = 1.11e-15$ |
